# Influence of flowering phenology and strobili amount on mating patterns in an *Abies nordmanniana* clonal seed orchard

**DOI:** 10.64898/2026.09.09.750367

**Authors:** Ole K. Hansen, Jing Xu, Ulrik B. Nielsen

## Abstract

Clonal seed orchards (CSOs) provide improved plant material from forest tree breeding. Mating dynamics of CSOs is therefore interesting, since dysfunctions like selfing, pollen contamination and differences in clonal contributions of gametes may hamper the genetic gain. Skewed paternal contribution can be caused by clonal variation in amount of flowering/pollen production, and clonal differences in flowering phenology. Including trees from different populations in CSOs entails a particular risk that the parents are split into non-overlapping phenological classes with limited gene exchange, so the seed crop will consist of different gene pools.

The study objective was to study the influence of flowering amount and flowering phenology on the mating patterns in an *Abies nordmanniana* CSO combining classic ocular observations of strobili with paternity analysis using DNA markers. Furthermore, we tested whether the two different gene pools from which the trees originated, had limited gene exchange.

Phenology observations over three years showed that female strobili in *A. nordmanniana* often are receptive before the male strobili release pollen (5-7 days). However, despite clonal variation in first pollen release there is a “great dusting day”, where all clones are contributing pollen. Based on DNA-markers, pollination success among 93 clones varied from 0 to over 5%. Clone variation in male strobili amount explained 29% of the variation in pollination success, while early pollen shedding could explain 13%. Our results support the hypothesis of early pollen shedding having an advantage in pollination success and late pollen shedding a disadvantage. No substantial mating barriers between trees from the two gene pools which are included in the CSO could be observed.

## Introduction

Seed orchards are the major tool to provide genetically improved plant material from most breeding programmes of forest trees, and clonal seed orchards (CSOs) is the most common type of seed orchard used for this task (White et al. 2007). It is therefore of major interest to study the mating dynamics of CSOs, since dysfunctions like selfing, pollen contamination from surrounding trees and skewed parentage due to differences in clonal contributions of gametes can severely hamper the genetic gain and make serious deviations from the panmictic equilibrium that is the assumption in an idealized seed orchard (Eriksson et al. 1973). A skewed paternal contribution can be caused by a straightforward clonal variation in amount of flowering and pollen production, but random mating may also be obstructed due to differences among clones in flowering phenology (Eriksson et al. 1973), where asynchronously flowering trees may not find mates. In CSOs, trees from different natural populations can be brought together. In combination with the strong genetic control of some components of flowering phenology, there is the risk that the CSO parents are split into non-overlapping phenological classes with limited gene exchange (Funda & El-Kassaby 2012). Therefore, if the parents in the CSO are coming from different gene pools, there is a risk that also the seed crop will consist of different gene pools. Evidently, variation in amount of flowering as well as its timing can influence the mating pattern at the same time, thereby making the use of any of the two factors as predictors for siring success more complicated.

In Denmark conifers of the genus *Abies* are used as exotic species in the forestry and in the Christmas tree and greenery production. For both types of production, CSOs have been delivering seeds for decades. Studies of mating patterns in Danish *Abies* seed orchards with DNA-markers have revealed the well-known phenomenon of skewed parentage, but also that pollen contamination is limited; several studies have shown it to be in the range of 1.4% to 6% (Hansen & Kjær 2006; Hansen 2008; Hansen & McKinney 2010; Hansen & Nielsen 2010; Xu et al. 2018; Hansen et al. 2020). This low contamination rate gives the possibility to study mating patterns in a relatively ‘undisturbed’ system where replicates of different genotypes also make it possible to make more basal studies of the mating patterns of conifers.

The Danish breeding programme for Nordmann fir Christmas trees consists of several gene pools. The main part of the breeding population originates from central Georgia in Caucasus, i.e. the Ambrolauri and Borshomi areas. All is based on plus-tree selection in Danish stands, and in the case of Borshomi both 1^st^ and 2^nd^ generation material. The CSO FP.272 Skelhusmarken is among the largest Danish Nordmann fir seed orchards and originally consisted of 100 clones comprising 75 plus-trees of 2^nd^ and 25 plus-trees of 1^st^ generation Danish material of Borshomi origin. Later, results from two provenance tests of the same material of origin have shown a remarkably change in vegetative budburst from 1^st^ to 2^nd^ generation, with the later having a budburst delayed up to 6 days (Nielsen et al. 2010), or 50% versus 20% of the trees, respectively, had initiated growth in early June (Nielsen 2006). If a similar difference was to be found in the flowering phenology, there is a high risk of asynchronously flowering. Christmas trees are a high value crop and high genetic quality of the seed used is important for economic success in a strongly competitive market. Therefore, estimating the genetic worth (Stoehr et al. 2004) based on an ocular strobili evaluation has been proposed, as a tool to declare the quality of the seed crops (Nielsen & Hansen 2012), but asynchrony may bias the estimated genetic worth to an unknown extend.

### Objectives of the study

As a general objective, we wanted to study the influence of both flowering amount and flowering phenology on the mating patterns in an *A. nordmanniana* CSO combining classic ocular observations and actual paternity analysis using DNA markers. As a more specific objective, we wanted to see whether the two different gene pools from which the trees originated, had limited gene exchange.

## Material and methods

### Clonal seed orchard - Plant material

In the CSO FP.272, situated at Himmerland State Forest district, 100 plus-trees were high-grafted in 1997 onto rootstocks of an existing *A. nordmanniana* plantation with a spacing of 5 x 5 m. The CSO covers an area of around 10 hectares. The 100 plus-trees are all considered to originate from the Borshomi region in Georgia, Caucasus, which is part of *A. nordmanniana*’s natural distribution area. The seed were imported around 1900, and gave rise to some stands, of which several later has been certified as seed stands (Løfting 1973). Two of these stands were established in Tversted and still exist, namely F.526 and F.527, where respectively 4 and 21 of the original 100 clones in FP.272 have been selected. These 25 clones are designated the “Tversted pool” in FP.272 and is 1^st^ generation Danish material of Borshomi origin. Another certified seed stand coming from the early seed import around 1900, and presumably of Borshomi origin, was established at Boller forest district and named F.20. It does not exist today, but has been the source for many later stands, including three stands from which the remaining 75 original clones in FP.272, the “Boller pool” (2^nd^ generation Danish material of Borshomi origin), were selected. As described earlier: although the material in both pools are thought to originate from Borshomi in Georgia, field trials have shown that offspring from the Boller clones in general have later flushing than offspring from the Tversted clones (Nielsen et al. 2010). Late flushing is very important in relation to Christmas tree production due to the risk of late spring frost, which damages the newly flushed shoots and makes the trees difficult/impossible to sell.

In the winter 2008-2009, seven of the 100 original clones in FP.272 were thinned away, based on assessment of post-harvest quality (unpublished data). Two were from the Tversted pool and the remaining five were from the Boller pool. For gene bank purposes, however, a single ramet of each of the seven culled clones was kept in the most western row of the CSO. So, in spring and autumn 2009, where the first strobili registrations and the seed harvest was conducted, FP. 272 contained 93 clones – each represented with around 33 ramets (mean 32.6, standard deviation = 1.2) – plus seven clones represented by only one ramet each. During the winter 2009-2010, an additional thinning was conducted, taking away 29 more clones, leaving 64 for the flower registrations in 2010 and 2011 (22 clones from the Tversted pool and 42 from Boller).

### Registration of timing and amount of flowering

Registrations of timing of flowering were done on a subset of the ramets in the seed orchard at regular interval throughout the flowering period. In 2009, registrations were done on seven dates: April 23, 26, 28, and 30 and May 2, 5 and 12. In 2010: May 11, 20 and 25. In 2011: April 29, May 1 and 4. A total of 488 trees were included in the registrations in 2009 (average = 5.2 ramets per clone; range 3 to 7 – depending on the distribution of ramets in the rows included in the two observational transects – see below). In 2010 and 2011, after thinning down to 64 clones, 372 trees (average = 5.8 ramets per clone; range 3 to 10) and 370 trees (average = 5.8 ramets per clone; range 4 to 10) were included in the registrations.

The registrations of the female cones, which are sitting in the very top of the trees, were conducted from a cherry picker (also known as a man lift or basket crane) that was “raised” to crown height along two transects (used as blocks in the statistical analyses, see below) at intervals that allowed for safe scoring of cone development stage by binoculars. The location of the two transects in the CSO is outlined in Figure 1. The following scale, inspired by Brøndbo (1971), was used to assess the timing/receptiveness of the female strobili: Score 1: from winter dormancy going to swollen bud. Tips of bract scales are not visible. Score 2: stage of bud burst. A ‘brush’ of tips of bract scales emerges from the bud. Score 3: bract scales are seen as up-right spikes, in overall appearance like a paintbrush. Score 4: the cone appears open. The bract scales tips range from an initially upright to a maximally horizontal position. Score 5: the cone appears more closed. All bract scale tips bend downwards or as minimum holds a horizontal position. Score 6: the cone appears closed. Bract scales shut totally or almost totally.

**Figure 1.**
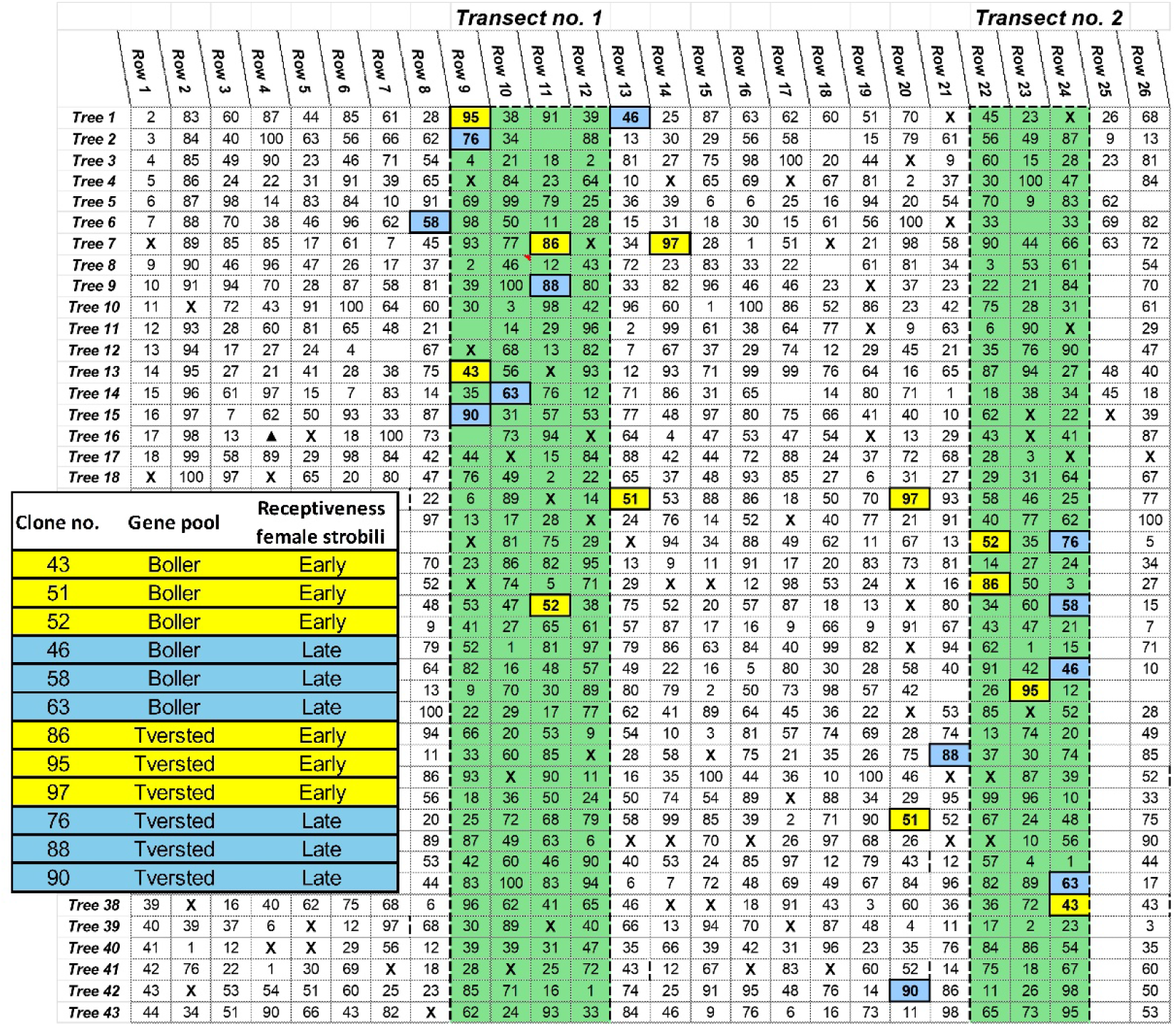
Sketch of a section of the CSO FP.272, where the two transects used for registration of flowering is marked with green (row 9-12 constitutes transect 1 and row 22-25 transect 2). Numbers are clone numbers. The southern part of the transects are not shown. The 24 seed catcher trees are also shown with yellow and blue colours – see text for details.

The registration of male strobili development stage was done on the same day immediately after finishing the scoring of the female cones. As male are sitting lower in the tree, the scoring was done from the ground, walking the same transects (Figure 1) and scoring each tree from its south-facing side. The following scale, which is a modified version of a scale developed by Webber & Painter (1996), was used to estimate how advanced the male strobili were in relation to pollen shedding: Score 1: the dormant winter bud. Score 2: the swollen meiotic buds in which the pollen grains are beginning to form. Score 3: buds are bursting their bud scales, but the microsporangia are still tightly packed, buds are still soaking wet if they are squeezed between the fingers. Score 4: strobili are mature and near shedding as microsporangia dries up, thereby loosening and separating into the mature structure. Score 5: strobili are shedding pollen, comprising three sub-categories: 5-1: beginning pollen shedding, 5-2: maximum pollen shedding and 5-3: almost finished shedding. Score 6: strobili are spend, dried-out brown structures.

Each ramet was assigned a lowest and highest value for both strobili scores, thereby describing the variation in receptivity and pollen shedding that may occur throughout the same tree. A binomial character was derived for cone receptivity (CONE), obtaining 1 as long as the ramet as a whole has a receptivity score of 4 or 5, otherwise 0. A similar character was made for pollen shedding (POLLEN), obtaining 1 as long any of the pollen shedding scores 5.1 to 5.3 was registered, otherwise 0.

The amount of male and female strobili in FP.272 was assessed by a logarithmic scale from 1 to 10 (score 1= 0 strobili, 2=1–3, 3=4–15, 4=16–60, 5=61–250, 6=251–1000, 7=1001–4000, 8=4001–16000, 9=16001-64000, 10= >64000) (modified from Sirikul et al. 1991). For male scores a maximum of 10, 8 and 9 was registered in the years 2009, 2010 and 2011 respectively and for female score a maximum of 6, 5 and 6.

### Vegetative phenology

Budburst was scored on one date in the years 2009 (May 19), 2010 (May 25) and 2011 (May 4) using a modified Langlet scale (Madsen 1994): score 0, buds in winter condition; score 1, buds start to swell, but not yet green; score 2, buds more or less green; score 3, bud burst; score 4, bud scales dropped, needles still turned forward like a brush; score 5, incipient shoot elongation, shoots and needles still soft and score 6, shoots elongated, needles in final position. Registration dates were chosen according to how progressed the flushing was in that particular year, aiming at getting the highest resolution in variation among the clones. A threshold value (BUDBURST) was calculated as the frequency of ramets within clone having a score>=3.

### Seed harvest, handling and germination

To pursue our goal of looking for differential pollination success, depending on the potential differences in flowering timing, twelve clones were chosen as pollen catchers in the CSO. These were selected based on preliminary analyses of the timing registrations conducted in the spring 2009. Six clones were chosen from each of the Tversted and Boller pools, and within the two pools, three clones had a relatively early receptiveness of the female strobili while the other three had a relatively late receptiveness (Table 1). Each clone was represented by two ramets (one in each transect) summing to 24 pollen catcher trees (Figure 1). Selection of the early receptive clones was based on early receptiveness as well as an early decrease in receptiveness. Selection of the late receptive pollen catcher clones was done based on clone values of the percentage of ramets showing no receptivity April 28 and 30 in 2009, the dates which gave the best resolution of the latest clones. The clones were ranked based on least square means data for female receptivity on the 28th. Additionally, information from the 30^th^ was used in choosing the latest clones by omitting those reaching 100% receptivity the 30^th^. Slightly different patterns for the two provenance pools were seen. The Tversted pool was very variable already on April 23, and we picked the three earliest at that date, which also tended to tail off early (May 5). The Boller pool showed a good distinction of clone receptivity April 26, and the earliest clones (100% on April 26) which also showed an early tail off in receptivity already on April 30 was chosen.

**Table 1.** Table showing the twelve clones chosen as pollen catchers in FP.272 by evaluating their overall pattern of female strobili receptiveness.

| Clone no. | Gene pool | Receptiveness female strobili |
| --- | --- | --- |
| 43 | Boller | Early |
| 51 | Boller | Early |
| 52 | Boller | Early |
| 46 | Boller | Late |
| 58 | Boller | Late |
| 63 | Boller | Late |
| 86 | Tversted | Early |
| 95 | Tversted | Early |
| 97 | Tversted | Early |
| 76 | Tversted | Late |
| 88 | Tversted | Late |
| 90 | Tversted | Late |

On September 17, 2009, cones were harvested from the two transects in the CSO; at each transect from one ramet of the twelve clones. Around 5-6 cones from each ramet were harvested and put into separate nets. After harvesting, the nets with cones were moved to a barn and hung on wires to secure air movement around the nets, and kept at approximately 10° C, until the end of October, then transferred indoors to a table of hard-cloth-wire to disintegrate at 20° C for about one week. Afterwards, the seeds were extracted manually, dried, cleaned and kept at 5° C in sealed plastic bags until the middle of November (20/11) where the seeds were put to storage at minus 6-7° C until stratification and germination. Seed stratification was initiated in the beginning of April 2010 (8/4), keeping each of the 2 x 12 = 24 seed lots separate. After soaking in running water for 24 hours, seed was kept refrigerated at 5° C in specially manufactured germination boxes based on a plastic box from Ultraplast (www.ultraplast.dk) type A6/60.

The first seeds started to germinate after around 9 weeks. Seeds showing emerging radicles were transferred to similar germination boxes at room temperature and typically grown for 2 weeks. The cotyledons and hypocotyl were then cut to pieces (around 50 mg) and used for later DNA-extraction. From each seed lot 48 seedlings were used, giving 1152 individuals.

In June 2010 needle samples from all 100 clones initially included in FP.272 (potential fathers) were collected for DNA extraction and genotyping. For the 7 + 29 = 36 clones where only one ramet remained in FP.272 after the genetic thinnings in 2008-2009 and 2009-2010, additional ramets were obtained from clonal archives established elsewhere. Each clone was thereby typically represented by two or, most frequently, three ramets.

### Laboratory work

All DNA extractions were done with the DNeasy Plant Mini Kit or DNeasy 96 Plant Kit from QIAGEN® (Germany). Genotyping with microsatellites was made in 10 μl PCR reactions using the QIAGEN® multiplex kit (catalogue no. 206143), following the given standard multiplex PCR protocol: 1 × Multiplex master mix (providing a final concentration of 3 mM MgCl2), 0.2 µM of each primer, around 20 ng of DNA sample and added water to make the final reaction volume. Amplifications took place in Perkin Elmer ABI thermal cyclers (model 2700 and 9700) with the following thermal profile: initially 15 min of denaturation at 95 °C, then 30 cycles of denaturation at 94 °C for 30 seconds, annealing at 58 °C for 90 seconds and extension at 72 °C for 60 seconds, with a final extension step at 60 °C for 30 min. One PCR reaction was made for each individual containing eight primer pairs originating from six different Abies species: NFH3, NFH15 and NFF3 (Hansen et al. 2005), Ab12 (Rasmussen et al. 2008), Abfi18 (Saito et al. 2005), SF b4 (Cremer et al. 2006), As09 (Lian et al. 2007) and AfSI_16 (Josserand et al. 2006). Fragment sizes of the amplified labelled microsatellites were determined on an ABI 3130XL genetic analyser and analysed with the GeneMapper software version 4.0 (Applied Biosystems).

### Data analysis

Statistical analyses of the field observations were carried out with the SAS ® software (SAS Institute Inc. 2013) using the GLM procedure for analysis of variance.

Model [1] was used for analyses within year and date for strobili counts and the binomial data for respectively cone receptivity and pollen shedding.

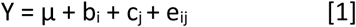

Where Y - observation, µ - overall mean, b – fixed block effect refers to the two transects used for data collection, c – fixed effect of clone and e - residual variance assumed independent and normally distributed. Least square means for clones were estimated and used for comparisons with results from paternity analysis.

Differences between the two gene pools were tested using the model [2]

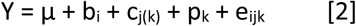

Where Y - observation, µ - overall mean, b – fixed block effect, c – fixed effect of clone within gene pool, p - effect of gene pool and e - residual variance assumed independent and normally distributed. Least square means for gene pools were estimated.

Model assumptions of normality of residuals and homogeneity of variance was checked by normality test and plotting residuals against predicted values using the plots=all option in SAS GLM. Analyses of scored values of strobili amounts benefits from the logarithmic nature of the used score, and assumptions were fulfilled to a satisfactory degree, which was also the case for the vegetative flushing score, based on an underlying continuous progress of the flushing. The frequency of ramets, within clone showing female receptivity or pollen shedding, deviated from the assumptions when frequencies were close to zero or one, but otherwise acceptable.

Pairwise correlations between traits were established using a simple linear model between the traits e.g. actual parentage as dependent and pollen quantity as independent variable using the SAS procedure GLM.

### Identity check in FP.272 trees

An initial allele frequency and identity analysis of the genotypic data from the around 280 trees sampled from FP.272 was performed using the software CERVUS 3.0 (Marshall et al. 1998; Kalinowski et al. 2007).

The aim was to establish how many genotypes were present in the seed orchard, which could be potential parents to the seedlings.

### Paternity analysis

For the seedlings from the 24 seed lots harvested in FP.272, a paternity analysis was conducted using the software CERVUS 3.0 (Marshall et al. 1998; Kalinowski et al. 2007) to identify paternity (mother known) for all seedlings. A list of candidate parents comprising all genotypes found in trees of FP.272 (see above) was used as potential parent information. Progenies with no paternal fit from the seed orchard clones were assumed to represent pollination from outside trees (pollen contamination).

The relative contribution of each clone to the gene pool of male gametes (pi) was calculated from the paternity analysis. The effect of low number of clones and/or unbalanced male contribution was quantified by the status number (NS) (Lindgren & Mullin 1998), which reflects build-up of coancestry in the seed orchard crop due to low number of clones and unequal male contribution (equation [1]):

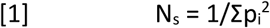

where Σp ^2^ is the sum of the squared male contributions (p ).

## Results

### Strobili counts and phenology

Large year-to-year variation was seen in strobili setting, where 2009 produced a very large crop of both male and female strobili, while the two following years were more moderate (Figure 2). The Tversted pool had more male strobili than the Boller pool, especially in 2009 (p<0.001), but did also differ in 2011 (p=0.004). The female strobili amounts were very similar between the two pools across years but was significantly higher in the Tversted pool in 2010 (p=0.003).

The clonal female strobili contribution was remarkably even in the large seed year 2009, where all clones contributed to the strobili crop (Figure 3). The two following years, having a relative female strobilus setting of less than 40% of 2009, still showed a rather even contribution, and no single clone added more than 7% of the female contribution in any year. For male strobili production, most clones had male strobili in 2009, almost all had in 2010, while the contribution was more skewed towards fewer clones contributing in 2011 (Figure 3).

**Figure 2.**
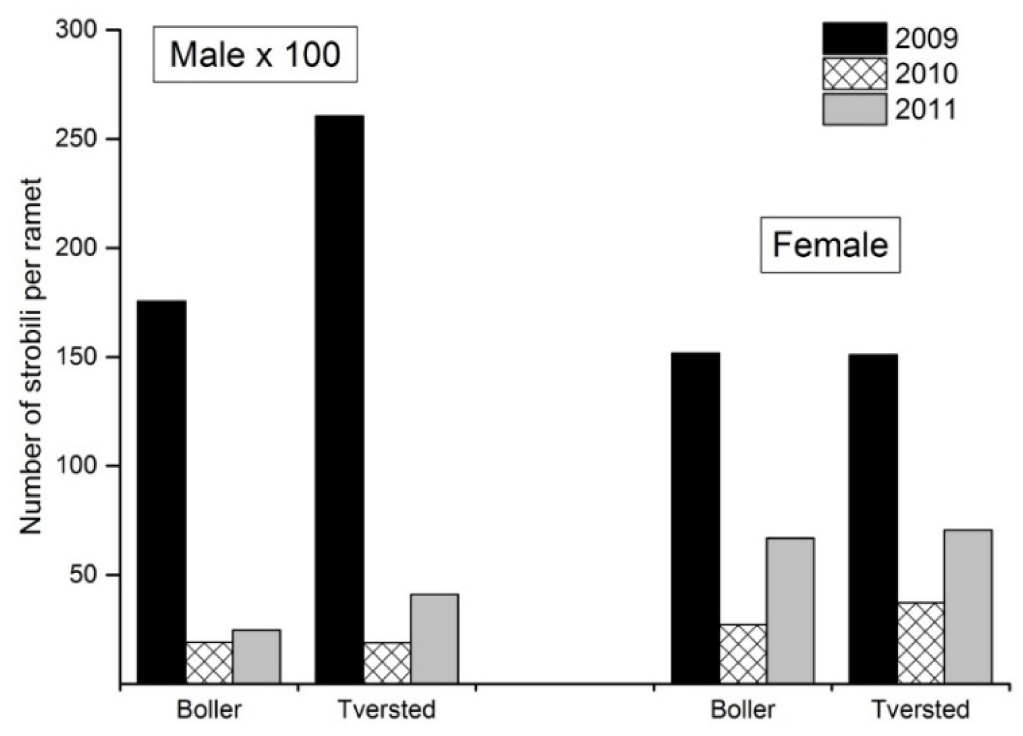
Overall number of male and female strobili per ramet in the three years of flowering registrations.

**Figure 3.**
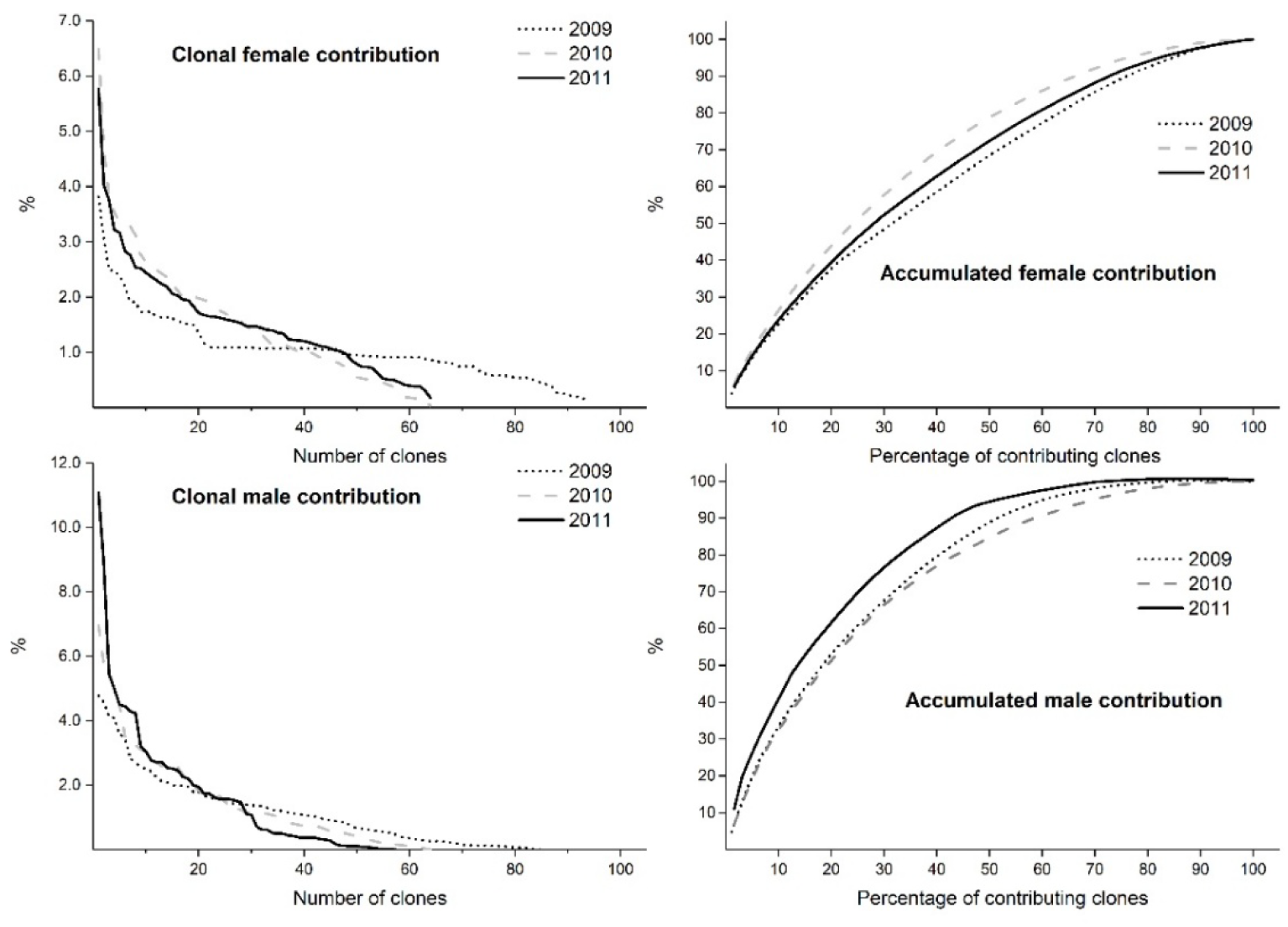
Clonal female and male strobili contribution (in %) sorted by descending contribution (left) and accumulated clonal strobili contribution (in %) against percentage of contributing clones (right).

In year 2009 more than 50% of the clones had receptive female strobili before the first pollen was shed in the orchard, however, all clones were shedding pollen and being receptive at the peak of the pollen release in 2009 on May 2 (Figure 4). Despite fewer registration dates, early receptiveness of females was also seen in 2010, but not all pollen-shedding clones were active on the same registration day. In 2011, we saw a similar picture as in 2009 - all clones with strobili being active at the peak day on May 1 (Figure 4). Showing the 2009 strobili data for the two gene pools separately (Figure 5), indicate a similar starting point for female receptivity and pollen shedding, but that the Boller pool had slightly more clones receptive for a couple of days (April 26 and 28) as well as pollen shedding (April 28 and 30). However, at peak-day (May 2) no differences were seen between the two gene pools (Figure 5).

The 12 clones picked as representative pollen catchers in 2009 were grouped in batches of three clones, i.e. early and late receptivity for the two gene pools. The expected differences in percent of ramets within clones scored as receptive were seen at the first three days of evaluation in 2009 – close to 100 % for the early clones versus no receptive clones at all for the late clones (Figure 6).

**Figure 4.**
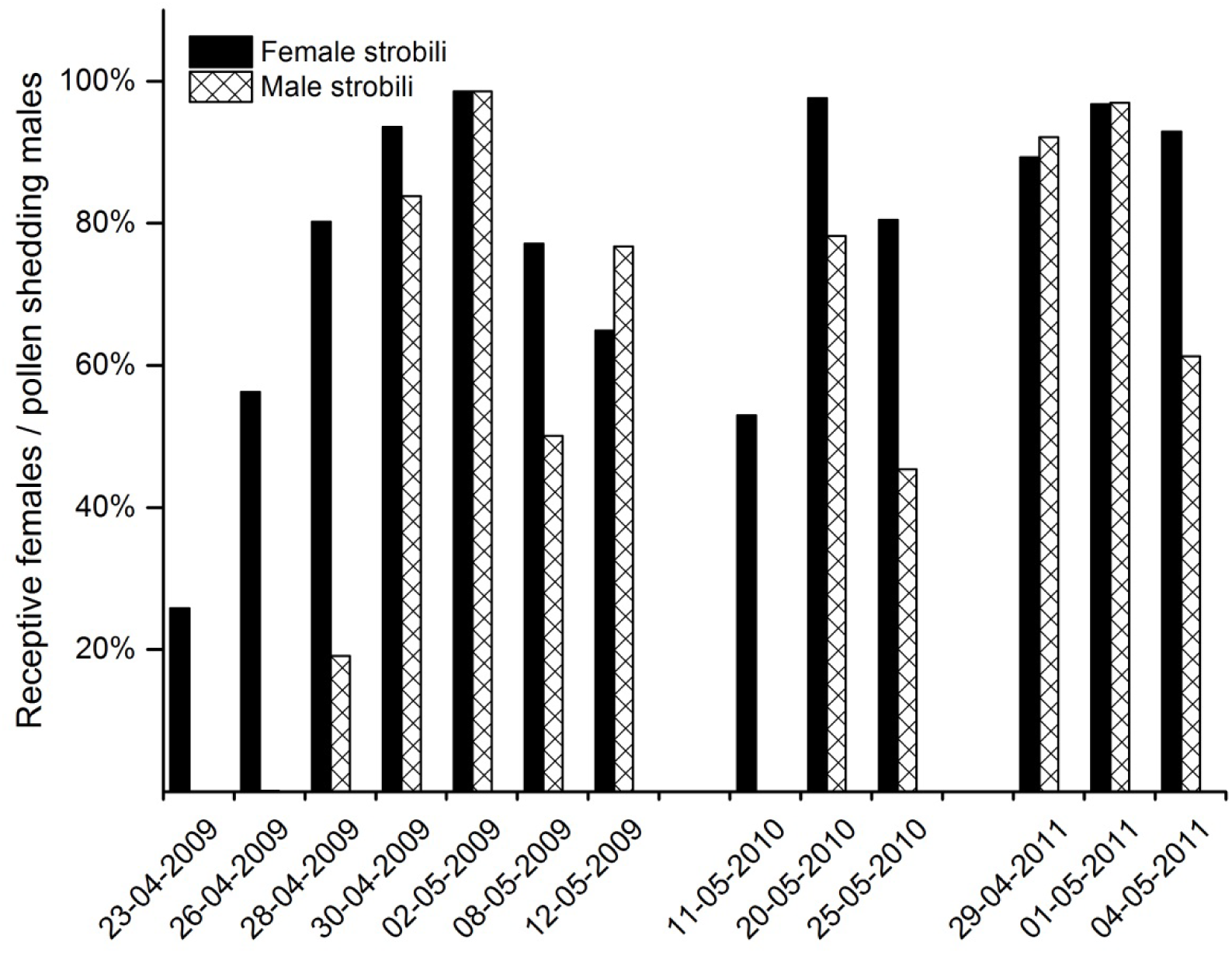
Frequency of receptive females and pollen shedding males on the registration dates during the three years.

**Figure 5.**
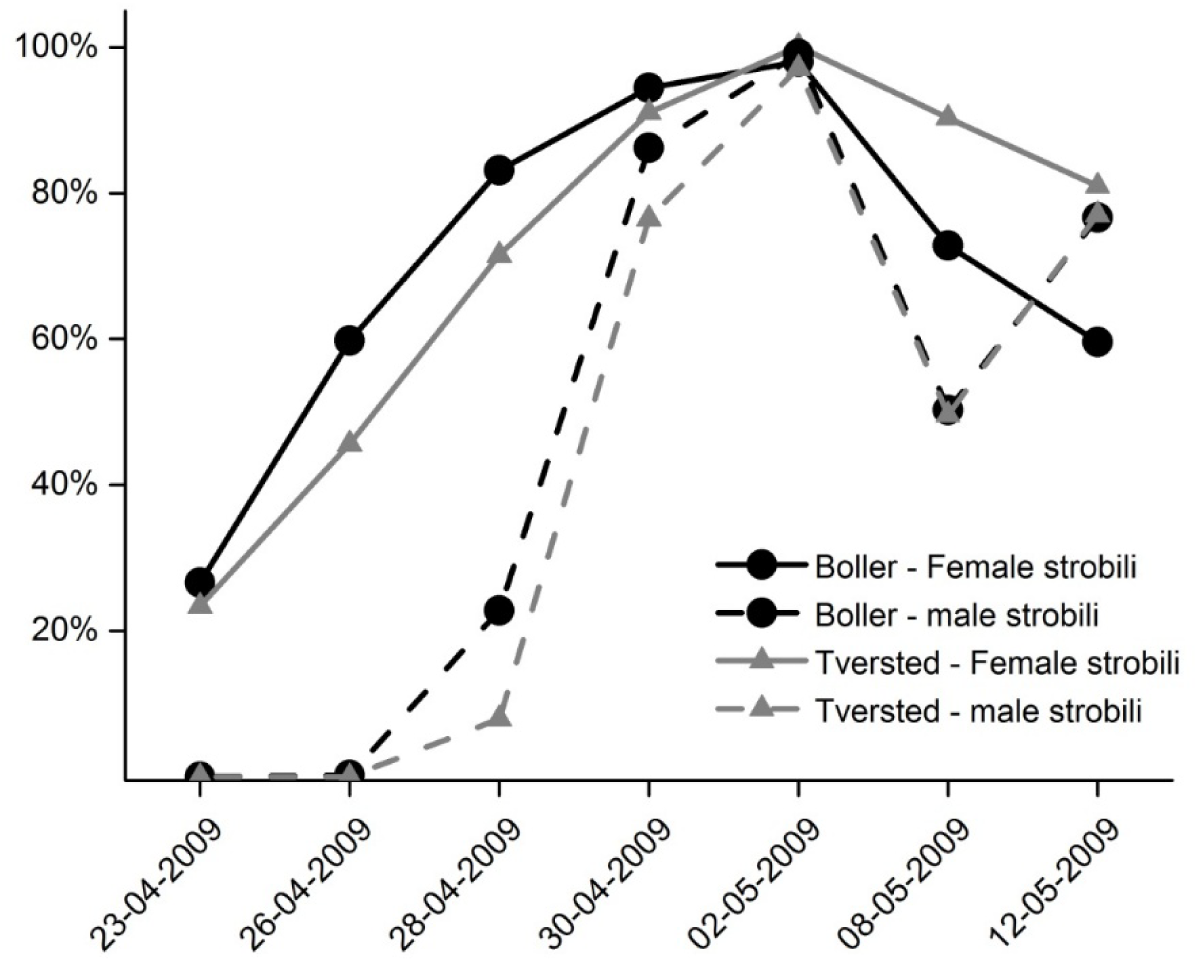
Timing of flowering in 2009, female receptivity (%) (solid line) and pollen shedding (%) (dashed line) across registration dates for the two different gene pools Boller (cirkel) and Tversted (triangle) .

**Figure 6.**
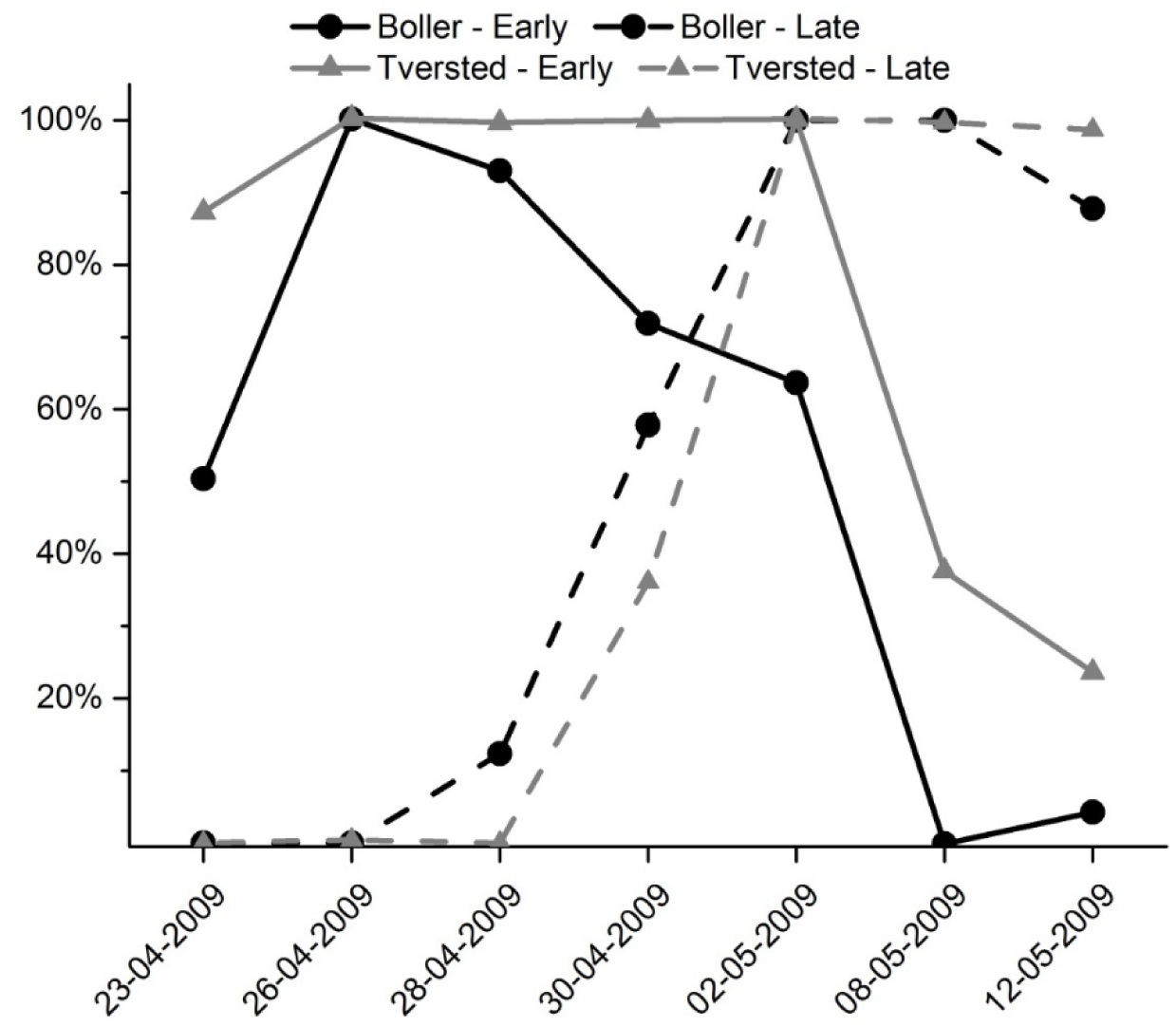
Timing of female receptivity (percentage) of the 12 pollen catchers for the two gene pools Boller (circle) and Tversted (triangle), respectively. Mean of the picked three early clones (solid) and three late clones (dashed) within gene pools.

The vegetative budburst was evaluated in all three years, and very consistent results were seen, where gene pools (p<0.001) and clones within gene pools (p<0.001) were significantly different. The Boller pool clones were latest flushing in all three years, and the percentage of trees showing budburst (score >=3) at the registration date was 69.9%, 91.7% and 54.0 % respectively in years 2009 to 2011. In comparison, the Tversted pool had 92.6%, 99.4% and 81.6 % for the corresponding years.

The difference in vegetative budburst between the two gene pools, also demonstrated in previous studies, was a major reason to our worries about two gene pools with no or little overlapping flowering. We therefore estimated the correlation between the vegetative phenology score and timing of respectively female receptivity (the binomial character CONE) and pollen shedding (the binomial character POLLEN). We focused on the year 2009, with most registration dates, and in 2009, we chose the date with the largest variation span for each of the two variables – April 26 for female cone receptivity and April 28 for pollen shedding. On a clonal mean level for all 93 clones in FP.272, there was a significant correlation of 0.25 (p=0.015; N=93) between vegetative phenology and female cone receptivity on April 26. Between vegetative phenology and pollen shedding on April 28 the correlation of 0.16 was not significant (p=0.134; N=93). Making this calculation separately for the two gene pools, gives a significant correlation between vegetative phenology and both female cone receptivity (r=0.39; p= 0.001) and pollen shedding (r=0.24; p= 0.047) for the 70 Boller clones. For the 23 Tversted clones, no correlation was found between vegetative phenology and female cone receptivity (r=-0.01; p= 0.970) nor pollen shedding (r= 0.28; p=0.201).

### Identity check and paternity analysis

Including the seven clones where only one ramet existed (culled in the winter before pollination), 104 genotypes were found among the genotyped ramets of the seed orchard. This number is due to four instances where two genotypes were found for trees with the same clone number. Typically, one genotype is correct and found in the two of the three genotyped ramets and the other is incorrect – called respectively A and B in Figure 7. The incorrect genotypes may for example be the result of a rootstock having overtaken the graft and becoming the stem in the tree.

Of the original 1152 seedlings, valid genotypic data was obtained for 1116 individuals – 36 individuals had too bad DNA quality to be satisfactory genotyped. Of these 1116 remaining seedlings, a maximum of 27 (=2.5 %) of the seedlings were found to be the result of incoming pollen (pollen contamination). Four seedlings were assigned two different fathers (with equal likelihood) and were omitted from the analysis.

**Figure 7.**
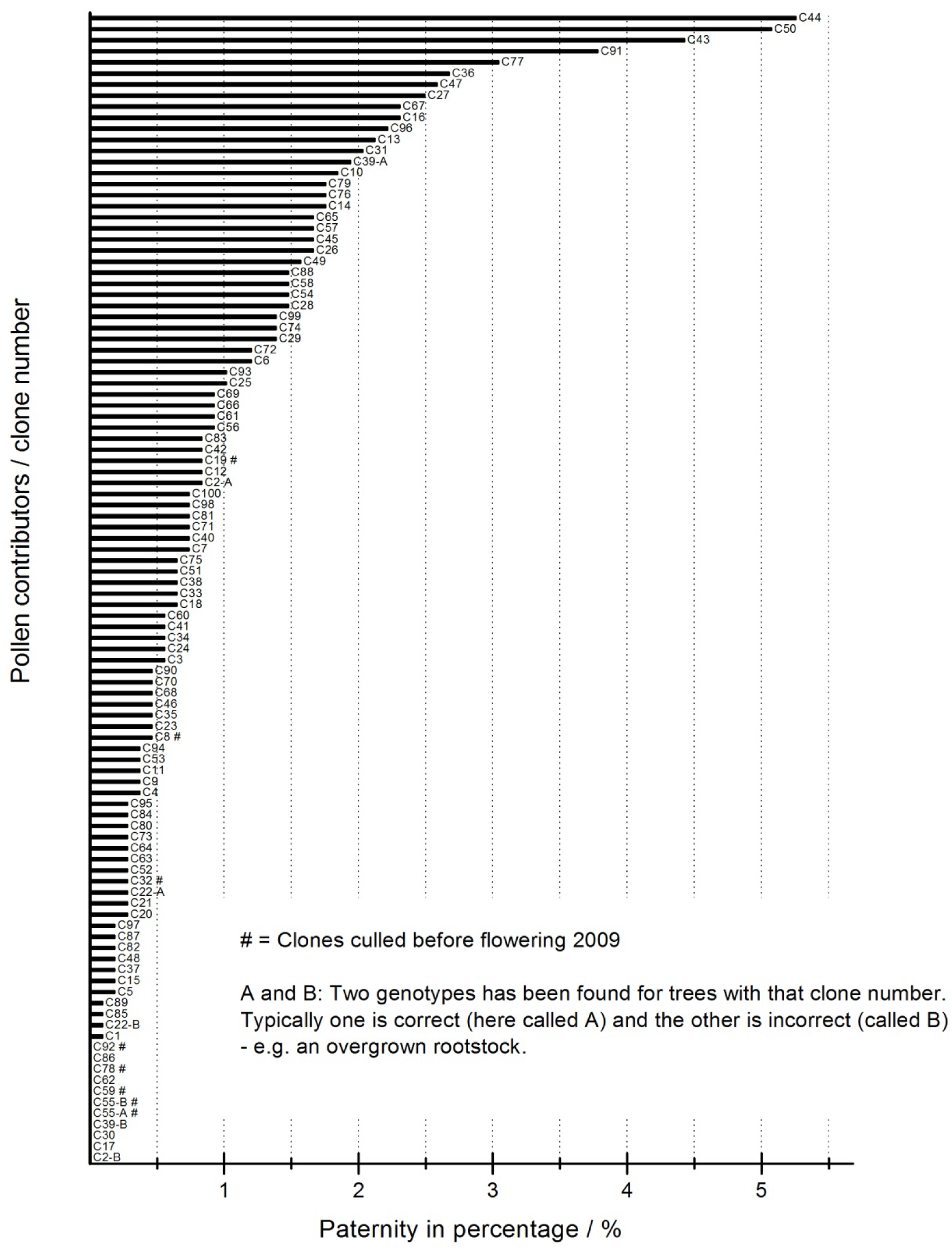
Distribution of paternity for 1085 genotyped offspring from FP.272. The pollen contributors marked with # are clones culled from the CSO, and where only one ramet should be present. Some clone numbers are represented twice (marked A or B) – these are clones where several genotypes where found.

Consequently, there were 1085 individuals to show the pollination pattern within the FP.272, and the distribution of pollination success for the 104 potential fathers is illustrated in Figure 7. It shows that pollination success varies a lot among clones – ranging from 0 % (11 genotypes) to a couple of clones having more than 5 % pollination success (C44 and C50). Since the ramet number of most clones was very balanced (mean 32.6, standard deviation = 1.2, range 27-34), the difference in pollination success was not due to varying number of pollinator trees. Several of the clones with no pollination success belong to the group of 7 clones which were culled in the winter 2008/2009 – i.e. clones C55, C59, C78 and C92 - an expected result due to the almost total removal of these genotypes from the CSO. On the other hand, three of the removed clones (C32, C8 and C19) are assigned paternity for respectively 3, 5 and 9 seedlings.

Especially the higher of these numbers indicate that the thinning may have left more than one ramet of some of the culled clones. Among the 11 genotypes with no pollination is also found 3 of the 4 genotypes which were assumed to be incorrect (called B) – and the third (C22-B) is only assigned paternity to one seedling. This ends up with, that only four clones (C17, C30, C62 and C86) of the 93 clones, which were intended to be pollinators, did not sire any of the sampled seedlings. Interestingly, these four clones were all in the third (roughly) of the clones which had the latest pollen release (based on the date 28/4-2009); i.e. their ranking was in the range of 61-91 out of the 93 clones in relation to early pollen release. The status number (N_S_) for the above-mentioned 93 clones, disregarding the minimal pollen contamination and the small/lacking contribution from the additional 11 discovered genotypes, was 47.

For the 12 clones acting as pollen catchers, an estimate for self-pollination could be obtained. These clones had together a pollination success of 11.7 % (127 seedlings). Of those 127 seedlings, 19 were the result of self-pollination – giving an overall selfing rate of around 15 %. So, for these particular clones, we have a higher frequency of selfers than could be expected based on the overall pollination success of the pollen catchers. However, this high number is caused by some of the clones having unusual high selfing. Eight of the 16 seedlings sired by clone C58, for example, was the result of mating with its own genotype.

### Combination of strobili and paternity results

The pollination success in relation to flowering amount was visualised by plotting the results from the paternity analysis against the clonal male strobili contributions (Figure 8). Although there is a clear relationship, there are also several deviating clones – for example C50 and C43 (Figure 8), which have a medium proportion of male strobili (∼1-1.4%) but around 4.6 - 5% of the pollinations. Remarkably, clones C50 and C43 are ranked as number 4 and 8 out of the 93 clones when it comes to early release of pollen (based on data from28/4-2009) and C44 (ranked 6) and C27 (ranked 30) were also early pollen releasers in 2009. Looking at the clones which showed the opposite pattern – i.e. having a relatively large share of the male strobili, but a kind of underperformance when it came to percentage of pollinations, there was here a tendency that these clones had late release of pollen. In example were clones C54, C93 and C77 ranked 63, 76 and 87, respectively. Not all complied with this pattern though; C36 and C88 were rather early pollen releasers (ranked 13 and 25, respectively) – even their pollination success was lower than could be expected from their share of male strobili (Figure 8). Overall, the proportion of contributed male strobili could only explain around 29% of the variation in pollination success – and a contributing factor to this low percentage is surely the influence of clonal variation in timing of flowering. The connection between earliness of male shedding and pollinations success was lower than pollen quantity, but still strongly significant (R²=13%, p<0.001). Combining clone male contribution and earliness into one single analysis can explain 35% of the actual pollination success.

**Figure 8.**
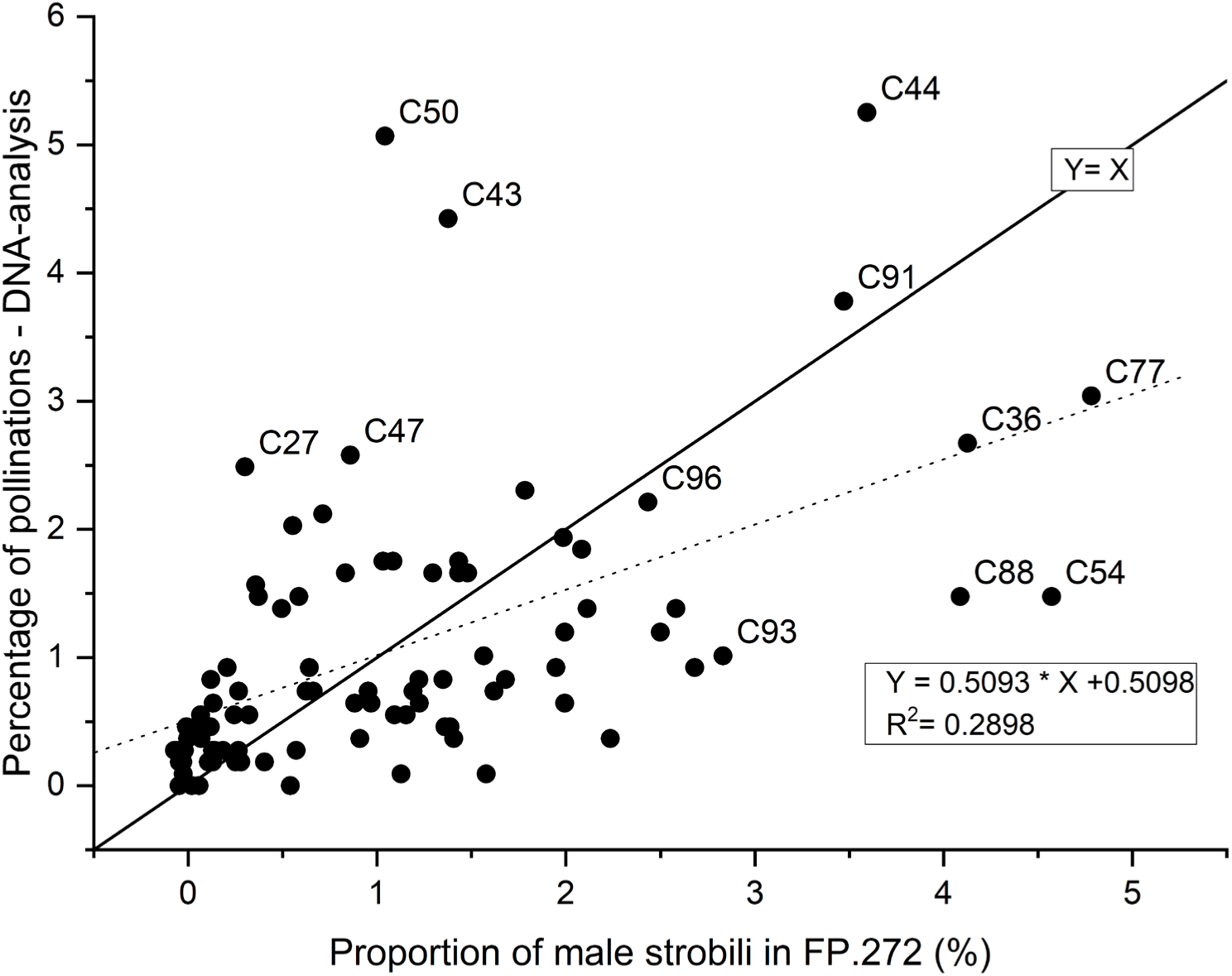
Relationship between clonal proportions of the total amount of male strobili in FP.272 in 2009 and the pollination success studied via DNA-markers. Dashed line shows the linear regression of pollinations against proportion of male flowers. Solid line is Y=X, which facilitates identification of distinct observations. There are only labels on clones positioned far apart from the rest in the plot.

For further elucidating the role of flowering phenology in pollination success, we looked for phenology patterns among the clones across gene pools. We identified three distinct groups concerning onset of pollen shedding on April 28 2009: 1) a group of 45 clones with late pollen shedding (only 0-4% of ramets had started shedding), 2) 34 clones in the middle (17-42% of ramets were shedding on the date) and 3) a group of 14 early clones (50-100% of ramets had started shedding) (Table 2). Looking at the number of pollinations the clones in the respective groups have made, it is seen that the 14 clones in the early shedding group (=15% of the clones), had made almost 29% of the pollinations (Table 2). Combined with the here observed tendency of Nordmann fir having receptive female strobili before pollen shedding trees in the CSO, the case seems evident: the early clones have a benefit of being early – they come first and are first served. However, if ones look at the amount of male strobili produced, the early group does also contribute disproportionally – where the 14 clones contributed with 25.3% of the strobili (Table 2).

However, the late shedding group has a disproportionate lower pollination success compared to estimated pollen contribution, namely a pollination success of 34.8% and male contribution of 41.1% supporting the importance of timing.

**Table 2.** Influence of timing of flowering on pollination success.

| Pollen release 28/4-2009 | # clones | % | # pollinations | % | Male strobili contribution % |
| --- | --- | --- | --- | --- | --- |
| Late clones (0-4%) | 45 | 48.4 | 371 | 34.8 | 41.1 |
| Middle clones (17-42%) | 34 | 36.6 | 390 | 36.6 | 33.5 |
| Early clones (50-100%) | 14 | 15.1 | 306 | 28.7 | 25.3 |
| Sum | 93 | 100 | 1067* | 100 | 100 |

\* Number is lower than the 1085 that are depicted in figure 7. This is because offspring sired by the clones, which were culled away, was omitted from this analysis, as no phenology data were available for these seven clones.

The distribution of pollinations for the two gene pools (Boller and Tversted) is seen in Table 3. Of the 1085 pollinations sired by members of the CSO, 848 (78.2 %) were originating from Boller clones while Tversted clones sired the remaining 237 (21.8 %). The estimated pollen contribution per tree was higher in the Tversted material, see Figure 2, leading to an estimated relative pollen contribution of 67.2 % for Boller and 32.8 % for Tversted, Table 3. On the other hand, the earlier pollen shedding in the Boller group, Figure 5, pulls in the other direction with regards to pollination success. The distribution of number of clones and ramets of the two pools was respectively 75.3 % / 24.7 % and 75.2 % / 24.8 % (Table 3), thereby being in rather good accordance with the pollination success.

**Table 3.** Distribution of pollinations etc. for the two gene pools.

|  | F.20 Boller | Tversted | Sum |
| --- | --- | --- | --- |
| # active clones | 70 | 23 | 93 |
| # active clones in % | 75.3 | 24.7 | 100 |
| # ramets | 2281 | 752 | 3033 |
| # ramets in % | 75.2 | 24.8 | 100 |
| # pollinations | 848 | 237 | 1085 |
| # pollinations in % | 78.2 | 21.8 | 100 |
| # male strobili – clonal average | 17,568 | 26,053 |  |
| Proportion of male strobili in %* | 67.2 | 32.8 | 100 |

\*Calculated as the average clonal value within the two gene pools multiplied with the number active clones, and then divided by the total sum of male strobili

Mating between individuals from different gene pools could be analysed from the pollinatoŕs point of view: when a clone made a successful pollination, how often did it mate with an individual from its own gene pool and how often with a tree from the opposite gene pool? The pollinator within each gene pool had a balanced opportunity to pollinate a Boller or a Tversted clone (6 pollen catcher clones from each gene pool, Table 1). Consequently, with no preferences, the expected pollination success would be 50% to pollen catchers from each of the two gene pools. For the Boller gene pool, 51.3 % of the 848 pollinations were made with another Boller clone (Table 4 upper part). For the Tversted gene pool, 52.7 % of the 237 pollinations were made with another Tversted clone – (Table 4 lower part). These numbers reveal a relatively higher frequency of within gene pool matings than between, but the difference is rather modest.

**Table 4.**
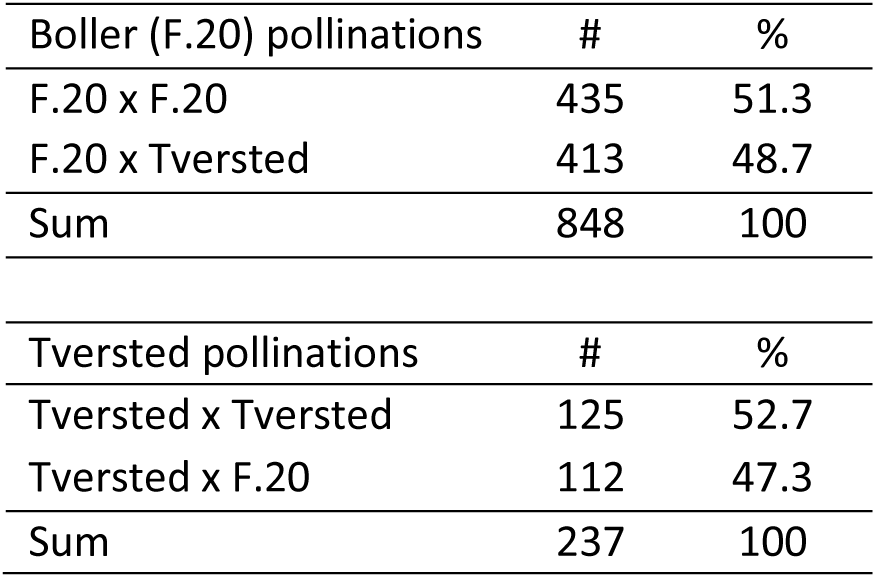
Mating within and between gene pools.

## Discussion

For a few, intensively breed conifer species, like *Pinus taeda* in the southern United States, large-scale full-sib family crossings are used in family forestry (McKeand et al. 2003). Vegetative propagation like somatic embryogenesis (SE) or rooted cuttings is also used in some conifers (e.g. *P. taeda* and *Picea abies*), but mainly to produce plant materials in breeding programmes and not for commercial scale. For SE this is a consequence of that the technique only works well with a few species (Pullman & Bucalo 2014). Thus, although the superiority of vegetative propagation methods and in particular SE has been pinpointed, the commercial use of SE is still limited (Lelu-Walter et al. 2013; Björs et al. 2025), and delivery of sexually reproduced seeds through seed orchards will continue to be the primary means of providing genetic improvement for the majority of tree species (Park & Bonga 2011).

In conifer CSOs, the skewed parentage, meaning that some clones contribute relatively more to the offspring than others, was documented through observational studies early in the use of CSOs, revealing great clonal variation in amount of flowering - a presumably good indicator for reproductive success (Sarvas 1962; Jonsson et al. 1976). On the female side, this was taken further, looking at clonal variation in cone weight or number of seeds (e.g. Griffin 1982; Schmidtling 1983) and showing similar results. On the male side, clonal pollination success has historically been notoriously more difficult to estimate than the clonal seed production. By the advent of allozymes in the early 1980’s it was possible to make studies of outcrossing versus selfing and estimates of pollen contamination (Adams & Joly 1980; Friedman & Adams 1985). A few allozyme studies were even able to make paternity test, although constrained by the limited polymorphism in these markers (e.g. Erickson & Adams 1989). However, it was first with the introduction of DNA-markers, and in particular highly polymorphic microsatellite markers, that detailed outcome of the mating patterns in the CSOs became feasible to obtain, and numerous have been made since – starting in the mid-2000’s (e.g. Moriguchi et al. 2004; Slavov et al. 2005; Hansen and Kjær 2006). Recently, detailed studies of mating patterns have also been conducted using single nucleotide polymorphism (SNP) markers (e.g. Bouffier et al. 2023), which are not so variable but obtain their discrimination power via a higher number of markers.

A few studies of conifer CSOs have tried to establish a relationship between amount of flowering (=reproductive investment) and actual reproductive success found by DNA-markers; e.g. in Japanese black pine (Goto et al. 2005), Sugi (Moriguchi et al. 2007), Nordmann fir (Hansen & Nielsen 2010) and lodgepole pine, Douglas-fir and western larch (Funda et al. 2011), and more recently in maritime pine (*Pinus pinaster*) (Bouffier et. al. 2023), and Chinese fir (*Cunninghamia lanceolata*) (Wu et. al. 2022) seed orchards.

The variation in flowering phenology has traditionally been assessed through morphological observations of different phenological classes (e.g. Brøndbo 1971), and asynchronous phenology has been observed in several conifer species such as Sitka spruce (El-Kassaby and Reynolds 1990) and Douglas-fir (El-Kassaby et al. 1984; Copes and Sniezko 1991). Contrary to the relative abundance case with observations and measures of the amount of flowering, studies that pair phenological observations in CSOs with detailed DNA-marker based paternity analysis to assess realized reproductive success have seldom been made. To our knowledge, the first study was the one of Sugi by Moriguchi et al. (2007), who made a simultaneously linear regression between male reproductive success and male strobili production and floral synchrony, with the latter two as independent variables. The two factors, however, only accounted for 15 % of the variation in male reproductive success. A similar approach in our study explained 35%. More recent studies in Chinese fir seed orchards have quantified clonal variation in flowering synchronization or, separately, assessed parental contributions through DNA-based paternity analyses, but these components have generally not been integrated to directly test how phenology and flowering intensity jointly determine genetic reproductive success (Xie et al. 2022; Wu et al. 2022).

In strobili amount, we found a substantial and significantly higher male production in the Tversted pool compared to the Boller pool in both 2009 and 2010. This may be due to the higher ontogenetic age of the Tversted clones; scions of which were taken in stands established around 1900. In comparison, the scions of the Boller clones were cut in three stands established in the period 1969-1979. As Nordmann fir normally starts to flower at the age of 30-35 years (Ousmael et al. 2023), some of the Boller clones were just entering their fertile age in 2009. In Nordmann fir, the female strobili, sitting in the very top of the trees, are normally observed some years before the male strobili that sit further down in the crown. This may explain why we saw no difference in female strobili production between the two gene pools.

Nordmann fir has periodically mast seeding years, so it was no surprise to see the large variation in strobili amounts across years, as also previously reported in Nordmann fir CSOs (Nielsen & Hansen 2012). In the bumper crop year 2009, the clonal contribution to the strobili production was remarkably evenly distributed. These results are, to some extent, backed up by the paternity analysis results, which showed that only four clones of the presumably 93 potential active clones did not sire any of the around 1100 genotyped seeds. The connection between number of male strobili and pollination success was definitely there, where male strobili contribution correlated positively with pollination success could explain 29 % of the variation in pollination success. The connection between earliness of male shedding and pollination success was lower than for pollen quantity, but still solely accounting for 13% in pollination success. In a previous study of a Nordmann fir CSO with 23 clones (Hansen & Nielsen 2010), the amount of male strobili could explain 76 % of the variance in pollination success. In that study information about flowering phenology was not included. However, the CSO studied in Hansen & Nielsen (2010) consisted of a relatively small number of clones where most of them came from the same region. With a higher number of clones, and originating from different gene pools, flowering phenology may have a significant impact on the mating patterns as well. This is pinpointed by some of the clones, which deviates most from the simple linear relationship between male strobili amount and pollination success (Figure 8), namely C43 and C50. These clones had only intermediate amounts of male strobili but were still among the most successful pollinators – which may be due to them being among the earliest pollen shedding clones – see further below.

However, despite the demonstrated variation in timing of flowering, the repetitive and detailed survey of the phenology in 2009 also showed, that there was what could be called ‘great pollen dusting day’, where all clones were shedding pollen and having receptive female strobili on May 2.

In two of the three study years, we saw a clear pattern of female strobili being receptive before pollen shedding. Franklin & Ritchie (1970), who studied cone phenology and shoot development of Noble fir (*Abies procera*) and other true firs in natural stands, reported the same phenomenon in Noble fir. A general earlier development of female strobili compared to male strobili was also reported in Norway spruce (*Picea abies*) (Nikkanen 2001). However, Nikkanen (2001) furthermore observed a single year where female and male flowering took place completely simultaneously, and the amount of pollen in the air was high right from the very beginning of the receptive period. This was attributed to warm weather and a resulting short flowering period, and this situation corresponds to our observations in 2011 (Figure 4), where April was the warmest ever measured in Denmark.

Our initial concern about having two gene pools in FP.272 with non-overlapping or only partial overlapping flowering, was based on already known distinct difference of budburst between the Tversted (1. generation) and Boller (2. generation) material. Overall, on a species level, there was no relationship between vegetative phenology and timing of flowering – illustrated by the fact that the late flushing Boller clones had earlier female receptivity and pollen shedding than the early flushing Tversted clones. However, on a clonal mean level we did find a significant correlation between vegetative phenology and female receptivity for all 93 clones in FP.272, while this was not the case for pollen shedding. Making the same calculations for the two gene pools separately showed a similar tendency for the 70 Boller clones, although now also with a significant correlation with pollen shedding. A tendency, which says that female strobili development is closer linked to vegetative flushing than male strobili pollen shedding.

Franklin & Ritchie (1970) described how cool, wet weather appeared to delay pollen shedding of fully developed male strobili in Noble fir, while female strobili growth did not seem to slow down during such weather conditions. Nikkanen (2001) also found that environmental factors had a stronger effect on male than on female phenology in Norway spruce, expressed by lower heritability for the former. These observations are thereby in line with the correlation patterns we found for Nordmann fir.

Our DNA-marker results confirmed that there could be a selective benefit of being an early pollen releaser – as these clones had a disproportionately large share of the pollinations and the late releasing clones had lower pollination success.

We found a pollen contamination rate of maximum 2.5% which emphasizes that this Nordmann fir seed orchard is highly suitable for pollination studies. This contamination rate is rather low compared to results from e.g. pine and spruce CSOs in other Nordic countries (Harju & Muona 1989; Pakkanen et al. 2000; Torimaru et al. 2009) – although see the later study by Funda et al. (2015). Low contamination rates in Danish *Abies* CSOs have been observed in a number of previous studies (see introduction for references) and are probably due to the modest timber production with *Abies* species in the forestry and that Christmas tree production take place in specialized production stands on a relatively small area where most trees are felled before sexual maturation.

## Conclusions

Our studies of flowering phenology in three consecutive years showed that female strobili in *Abies nordmanniana* often are receptive before the male strobili release pollen (5-7 days in 2009 and 2010). It also showed, that despite clonal variation in first pollen release there is a “great dusting day”, where all clones are contributing pollen. Paternity analysis revealed only a little pollen contamination (range 0.7 - 2.5 %). Based on DNA-markers, pollination success among the 93 potential paternal clones varied from 0 to more than 5%.

Clone variation in male strobili amount (pollen production) could explain 29% of the variation in pollination success. Similar, could early pollen shedding explain 13% of the variation in pollination success. Our results support the hypothesis of “first in first served” of early pollen shedding having an advantage in pollination success and late pollen shedding a disadvantage. No substantial mating barriers between trees from the two gene pools which are included in the CSO could be observed.

## Acknowledgments

We thank the Danish Nature Agency for giving access to the seed orchard and for lending us the man lift used for flowering registrations. Thanks to Lena Byrgesen for doing the many DNA extractions, to Ole Byrgesen for help in relation to seed handling and to Kirstine B. Nielsen for the huge work of cleaning and genotyping seeds and checking of data. Finally, we are indebted to Lars Nørgaard Hansen for the fine and thorough flowering registrations – often done on long working days and in a storm of pollen.

## Data availability

Data are available online: [PERSISTENT WEB LINK TO DATASETS].

